# Identification of sulfenylated cysteines in *Arabidopsis thaliana* proteins using a disulfide-linked peptide reporter

**DOI:** 10.1101/2020.03.25.989145

**Authors:** Bo Wei, Patrick Willems, Jingjing Huang, Caiping Tian, Jing Yang, Joris Messens, Frank Van Breusegem

## Abstract

In proteins, hydrogen peroxide (H_2_O_2_) reacts with redox-sensitive cysteines to form cysteine sulfenic acid, also known as *S*-sulfenylation. These cysteine oxidation events can steer diverse cellular processes by altering protein interactions, trafficking, conformation, and function. Previously, we had identified *S*-sulfenylated proteins by using a tagged proteinaceous probe based on the yeast AP-1–like (Yap1) transcription factor that specifically reacts with sulfenic acids and traps them through a mixed disulfide bond. However, the identity of the *S*-sulfenylated amino acid residues remained enigmatic. Here, we present a technological advancement to identify *in situ* sulfenylated cysteines directly by means of the transgenic Yap1 probe. In *Arabidopsis thaliana* cells, after an initial affinity purification and a tryptic digestion, we further enriched the mixed disulfide-linked peptides with an antibody targeting the YAP1C-derived peptide (C_598_SEIWDR) that entails the redox-active cysteine. Subsequent mass spectrometry analysis with pLink 2 identified 1,745 YAP1C cross-linked peptides, indicating sulfenylated cysteines in over 1,000 proteins. Approximately 55% of these YAP1C-linked cysteines had previously been reported as redox-sensitive cysteines (*S*-sulfenylation, *S*-nitrosylation, and reversibly oxidized cysteines). The presented methodology provides a noninvasive approach to identify sulfenylated cysteines in any species that can be genetically modified.

## INTRODUCTION

Biotic and abiotic stresses increase the production of reactive oxygen species (ROS) in plants. Hydrogen peroxide (H_2_O_2_) is recognized as a secondary messenger and can cause posttranslational modifications on proteins by oxidizing sulfur-containing amino acids, such as methionine and cysteine (Jacques et al., 2015;Waszczak et al., 2015). Cysteine is one of the least abundant amino acids, representing 1.86% of all amino acids in *Arabidopsis thaliana* (The Uniprot Consortium, 2019), but it is highly reactive and often functionally conserved in active protein sites (Marino and Gladyshev, 2010). Reaction of H_2_O_2_ with proteinacious redox-sensitive cysteine thiols leads to the formation of cysteine sulfenic acid (-SOH), a reaction that mainly depends on the *p*K_a_ of the cysteine and its local structural environment (Roos and Messens, 2011; Gupta and Carroll, 2014). Sulfenic acids are generally unstable and frequently an intermediary modification *en route* to more stable oxidation forms (Gupta and Carroll, 2014). For instance, -SOH can form intra- or intermolecular disulfides or mixed disulfides with another free thiol or glutathione (GSH), making it enzymatically reversible by the action of thioredoxins (Trxs) or glutaredoxins (Grxs), respectively (Roos and Messens, 2011; Akter et al., 2015b). Recently, extracellular H_2_O_2_ has been shown to be sensed through disulfide formation of extracellular cysteines in the plasma membrane receptor HYDROGEN PEROXIDE-INDUCED Ca^2+^ INCREASES1 (HPCA1), leading to Ca^2+^ influx in guard cells (Wu et al., 2020). Conversely, besides disulfide formation, -SOH can further oxidize towards sulfinic (-SO_2_H) and sulfonic acid (-SO_3_H). Whereas -SO_3_H is generally considered as an irreversible modification associated with protein degradation (Huang et al., 2018), -SO_2_H can be reduced via sulfiredoxins (Srxs) (Biteau et al., 2003; Akter et al., 2018). Protein *S*-sulfenylation can directly regulate protein functions. For instance, in *Arabidopsis*, H_2_O_2_-dependent *S*-sulfenylation of the BRASSINAZOLE-RESISTANT1 (BZR1) transcription factor promotes the interaction with other transcriptional regulators and drives gene expression (Tian et al., 2018). Alternatively, sulfenylation of catalytic cysteines can directly inhibit the enzymatic activity (Tanner et al., 2011; Gurrieri et al., 2019). As such, the identification of sulfenylated cysteine sites is a crucial step to advance our understanding of redox-regulated processes.

Over the past decade, indirect and direct approaches have been developed to capture and identify *S*-sulfenylated proteins (Takanishi et al., 2007; Yang et al., 2014; Gupta et al., 2017). Initially, carbon nucleophilic SOH-selective probes enabled the *in situ* detection of *S*-sulfenylation at the protein level in *Arabidopsis* and human cells (Leonard et al., 2009; Paulsen et al., 2012; Akter et al., 2015a). Further advancements in affinity-based enrichment strategies allowed the accurate identification of the sulfenylated cysteine residues within the proteins in both human and plant cells (Yang et al., 2014, 2015; Akter et al., 2018; Huang et al., 2019). In addition to these chemoproteomics approaches, a genetic construct based on the yeast (*Saccharomyces cerevisiae*) AP-1–like (Yap1) transcription factor was used to detect *S*-sulfenylated proteins. Yap1 forms mixed disulfides via its redox-active Cys598, located in the C-terminal cysteine-rich domain (cCRD), with the sulfenylated Cys36 of the oxidant receptor protein 1 (Orp1) (Delaunay et al., 2002). A Yap1-cCRD construct, in which Cys620 and Cys629 were mutated to Ala and Thr, respectively, and solely the redox-active Cys598 was retained, was used for the identification of *S*-sulfenylated proteins in *Escherichia coli* (Takanishi et al., 2007), yeast (Takanishi and Wood, 2011), and the legume model plant *Medicago truncatula* (Oger et al., 2012). In *Arabidopsis* cells, we generated a Yap1-cCRD construct fused to a tandem affinity purification (TAP) tag for improved capture and downstream identification of *S*-sulfenylated proteins (Waszczak et al., 2014). With this Yap1-cCRD construct, designated YAP1C hereafter, 97 and 132 *S*-sulfenylated proteins had previously been detected in the *Arabidopsis* cytosol and chloroplast, respectively (Waszczak et al., 2014; De Smet et al., 2019), but the sulfenylated cysteines remained unknown. Here, we describe how a tailored double affinity purification strategy enables the identification of *in situ* sulfenylated cysteines in a noninvasive manner.

## MATERIALS AND METHODS

### Plant materials and growth conditions

Transgenic cells expressing the YAP1C construct were generated as previously reported (Waszczak et al., 2014). In summary, the Yap1 C-terminal cysteine-rich domain (cCRD) construct, entailing the *Saccharomyces cerevisiae* Yap1-coding region corresponding to Asn565 to Asn650, was codon-optimized for expression in *Arabidopsis thaliana* (L.) Heynh. and synthesized with introduction of the mutations Cys620Ala and Cys629Thr. This genetic construct was fused with an N-terminal TAP tag, containing two IgG-binding domains of protein G and a streptavidin-binding peptide (SBP), separated by the Human Rhinovirus (HRV) 3C protease cleavage site. The YAP1C probe driven by a cauliflower mosaic virus 35S promoter was transformed in *Arabidopsis* cells. YAP1C expression levels were assessed by Western blot analysis (Waszczak et al., 2014). The PSB-D *Arabidopsis* cell suspension cultures (NASC stock no. CCL84840) were maintained as described in the ABRC Cell Culture Handling Protocol (https://abrc.osu.edu/researchers/cell). For H_2_O_2_ treatments, 500 mL of mid-log phase (3 days after culture refreshing, OD_600_=0.9) cells in 1-L glass flasks were treated with 20 mM H_2_O_2_ for 30 min before the cells were harvested through a vacuum filtration system (Pall Corporation, New York, N.Y., USA) and snap-frozen in liquid nitrogen before storage at −70°C.

### Protein extraction

Frozen cell pellets harvested from approximately 1 L of suspension cultures were crushed with quartz fine granules (Merck, Darmstadt, Germany) with a precooled mortar and pestle in ice-cold lysis buffer (25 mM Tris, 15 mM MgCl_2_, 150 mM NaCl, 15 mM pNO_2_PhenylPO_4_, 60 mM β-glycerophosphate, 0.1% NP-40, 0.1 mM Na_3_VO_4_, 1 mM NaF, 1 mM phenylmethylsulfonyl fluoride, 1 µM trans-epoxysuccinyl-L-leucylamido(4-guanidino)butane [E64], 1/50 mL ethylenediaminetetraacetic acid [EDTA]-free Ultra Complet tablet, 5% [v/v] ethylene glycol, 0.1 mg/mL 4-(2-aminoethyl) benzenesulfonyl fluoride hydrochloride [AEBSF], 0.1% benzonase, and 1 µg/mL leupeptin, pH 7.6), supplemented with 10 mM iodoacetamide [IAM] to prevent artificial oxidation of cysteine residues during the extraction procedure. After two rounds of centrifugation (20,000*g* for 20 min; 4°C), the supernatant was collected and protein concentrations were determined with the Bradford Protein Assay (He, 2011).

### Anti-C_598_SEIWDR antibody production and its coupling on magnetic beads

The C_598_SEIWDR peptide was synthetized (purity > 85%) and conjugated to Keyhole Limpet Hemocyanin (KLH) as a carrier (GenScript, Nanjing, China) and 0.2 mg of the C_598_SEIWDR-KLH conjugate, together with Freund’s incomplete adjuvant, were injected subcutaneously into four New Zealand rabbits at 14, 28, and 42 days. Seven days after the second and third immunization, approximately 20 and 40 mL (60 mL) of serum, respectively, were obtained from each animal. Three sera were retained for further purification, based on their high specificity against the ‘C_598_SEIWDR’ peptide (high ELISA titer, >1: 512,000), with a ‘C_598_SEIWDR’ peptide-coupled affinity iodoacetyl resin. Subsequently, the anti-C_598_SEIWDR antibodies were coupled on BcMag™ Epoxy-Activated Magnetic Beads (Bioclone Inc, San Diego, CA, USA) (Hamperl et al., 2014). Five mg of antibody was diluted to 3 mg/mL with coupling buffer (0.1 M sodium phosphate, pH 7.4) and incubated with 15 mg equilibrated Epoxy-Activated beads for 18 h at 30°C with gentle rotation (1000 rpm). After the beads had been washed twice with 100 mM glycine-HCl (pH 2.5) and then once with 10 mM Tris-HCl (pH 8.8), they were inactivated by 0.1 M trimethylamine and washed four times with phosphate-buffered saline (PBS) buffer (pH 7.4) and then twice with the PBS buffer (pH 7.4) containing 0.5% (w/v) Triton X-100. Finally, the antibody-coupled beads were suspended in 1 mL PBS buffer (pH 7.4), containing 0.02% (w/v) sodium azide and stored at 4°C until use.

### Affinity purification

Protein extracts (150 mg per sample) were incubated with 300 µl of IgG-Sepharose 6 Fast Flow beads (GE Healthcare, Chicago, IL, USA) for 2 h at 4°C with gentle rotation (1000 rpm). Then the beads were washed five times with washing buffer (10 mM Tris, 150 mM NaCl, 1 µM E64, 0.5 mM EDTA-free Ultra Complet tablet, 0.1 mg/mL AEBSF, and 1 µg/mL leupeptin, pH 7.6) and twice with digestion buffer (50 mM Tris-HCl buffer, pH 8.0). Hereafter, approximately 0.6 mg IgG-enriched proteins were first digested on the beads with mass spectrometry-grade Trypsin/Lys- C Mix (Promega, Madison, WI, USA) at a 1:50 (enzyme/substrate) ratio overnight at 37°C. Additional trypsin at a 1:100 (enzyme/substrate) ratio was added for an extra 4 h at 37°C. After the peptide-containing elution had been collected by gentle rotation n (1000 rpm for 1 min at 4°C), the beads were eluted twice again with digestion buffer. All three elutions were pooled. One-sixth of the tryptic digestion was used for protein-level identification with liquid chromatography-tandem mass spectrometry (LC-MS/MS) analysis. The remainder of the tryptic digestion was incubated with the magnetic beads coupled to 200 µL anti-C_598_SEIWDR antibodies for 2 h at 4°C with gentle rotation (1000 rpm). The tube was placed in the magnetic separator until the beads were captured on the magnet side, whereafter the clear supernatant was removed. Collected beads were washed three times with cold washing buffer (10 mM Tris, 150 mM NaCl, pH 7.6). The enriched peptides were eluted by incubation with 400 µL of 0.2 M glycine buffer (pH 2.5) for 10 min with rotation (1000 rpm). The supernatant was collected on the separator and supplemented with 100 µL 1 M Tris-HCl buffer (pH 9.0) for neutralization. After desalting with OMIX C18 pipette tips (Agilent, Santa Clara, CA, USA), the peptide samples were eluted with 100 µL 75% (v/v) acetonitrile containing 0.1% (v/v) formic acid and dried by vacuum centrifugation. The dried peptide samples were subjected to LC-MS/MS.

### LC-MS/MS

For the LC-MS/MS analyses, a Q Exactive Plus instrument was used (Thermo Fisher Scientific, Waltham, MA, USA) operated with an Easy-nLC1000 system (Thermo Fisher Scientific). Samples were reconstituted in 0.1% (v/v) formic acid, followed by centrifugation (16,000*g* for 10 min). The supernatants were pressure-loaded onto a 2-cm microcapillary precolumn packed with C18 (3 µm, 120 Å; SunChrom, Friedrichsdorf, Germany). The precolumn was connected to a 12-cm 150-µm-inner diameter microcapillary analytical column packed with C18 (1.9 μm, 120 Å; Dr. Maisch GmbH, Ammerbuch-Entringen, Germany) and equipped with a home-made electrospray emitter tip. The spray voltage was set to 2.0 kV and the heated capillary temperature to 320°C. The LC gradient A consisted of 0 min, 8% B; 14 min, 13% B; 51 min, 25% B; 68 min, 38% B; 69-75 min, 95% B [A, water; B, 80% (v/v) acetonitrile] at a flow rate of 600 nL/min. Higher-energy collisional dissociation (HCD) MS/MS spectra were recorded in the data-dependent mode with a Top20 method. The first MS spectra were measured with a resolution of 70,000, an AGC target of 3e^6^, a maximum injection time of 20 ms, and a mass range from *m/z* 300 to 1,400. The HCD MS/MS spectra were acquired with a resolution of 17,500, an AGC target of 1e^6^, a maximum injection time of 60 ms, a *m/z* 1.6 isolation window, and a normalized collision energy of 30. The *m/z* of the peptide precursors that triggered MS/MS scans were dynamically excluded from further MS/MS scans for 18 s.

### MS/MS data processing

RAW files were examined with the pLink 2 algorithm version 2.3.5 (Chen et al., 2019) and converted to MGF files by the MSFileReader (Thermo Fisher Scientific). Spectra were searched against representative Araport11 proteins (27,655 entries, v1.10.4, release on 06/2016), supplemented with the YAP1C protein sequence. Notably, the last seven amino acids ‘RDWLESC’ of AT1G74260, of which the reverse sequence is an isobaric ‘CSELWDR’ peptide, were omitted, because they resulted in high-scoring decoy matches, reflecting the enriched YAP1C-derived ‘C_598_SEIWDR’. A precursor tolerance of 20 ppm and a fragment mass tolerance of 20 ppm (HCD spectra) were specified. A specific tryptic search was used with a maximum of two allowed missed cleavages. Variable modifications included methionine oxidation, cysteine carbamidomethylation, and protein N-terminal acetylation. No fixed modifications were set. Results were filtered at a false discovery rate (FDR) threshold of 1% at the spectrum, peptide, and protein levels.

### Bioinformatics

#### Cross-linked peptide-to-protein assignment

Cross-linked peptides were extracted from pLink 2 cross-linked peptide reports (≤ 1% FDR). Due to trypsin miscleavages, redundant cross-linked peptides were identified for approximately 10% of the YAP1C cross-linked protein sites. To remove this redundancy, cross-linked peptides matching an identical protein site were grouped and the peptide matching the least proteins was chosen as representative, resulting in 1,748 YAP1C-cross-linked protein sites (Supplementary Dataset S1).

#### Enrichment analyses

The 570 proteins that uniquely matched cross-linked peptides were analyzed for gene set enrichment in the Gene Ontology (GO) and the Kyoto Encyclopedia of Genes and Genomes (KEGG) pathways by means of the DAVID tool (Huang et al., 2009). In addition, the overrepresentation of Cys-SOHs was determined in protein domains (PROSITE profiles) (Sigrist et al., 2013) as described previously (Huang et al., 2019).

#### C_598_SEIWDR cross-link peptide fingerprint scanning

Peak list (.MGF) files generated by pLink 2 (Chen et al., 2019) were parsed by an in-house script to detect the occurrence of the 10 characteristic fragment ions. Per MS/MS spectrum, the numbers of peaks with a matching *m/z* value (**≤** 0.01 Da) were counted, irrespective of their intensity. MS/MS spectra containing the full fingerprint or a high number of characteristic fragment ions hint at the fragmentation of CSEIWDR cross-linked (CL) peptides, although a CL search algorithm remains necessary to uncover their identity.

## Data availability

Thermo RAW files and pLink 2 result tables are available on the PRIDE repository with the identifier PXD016723. The 1,747 sulfenylated cysteine residues identified were submitted to the Plant PTM Viewer (Willems et al., 2019). Upon acceptance, the dataset will become publicly available in the Plant PTM Viewer.

## RESULTS AND DISCUSSION

### Identification of YAP1C-resolved *S*-sulfenylated sites through a protein- and peptide-level purification strategy

Previously, we had identified *S*-sulfenylated *Arabidopsis* proteins *in vivo* by means of a transgenically expressed YAP1C probe fused to an affinity purification tag (Waszczak et al., 2014; De Smet et al., 2019). Crucial steps in this strategy were the nucleophilic attack and the subsequent formation of a mixed disulfide bond by Cys598 of YAP1C and the sulfenylated Cys (Cys-SOH) residue in oxidized target proteins (Delaunay et al., 2002; Takanishi et al., 2007). In these approaches, the mixed disulfide YAP1C complexes were purified in a sequential affinity purification strategy, firstly by IgG-Sepharose beads trapping the IgG-binding domain of protein G (ProtG) and, subsequently, after cleavage with protease, by Streptavidin-Sepharose beads targeting the streptavidin-binding peptides (SBP). As such, approximately 230 *in vivo S*-sulfenylated protein targets had been detected by MS (Waszczak et al., 2014; De Smet et al., 2019). However, information on the identity of *S*-sulfenylated sites remained unknown in the proteins harboring more than one cysteine. An average of 7.85 Cys residues per *Arabidopsis* protein (The Uniprot Consortium, 2019) implicated that for most *S*-sulfenylation proteins downstream validation experiments are necessary, including mutational approaches and/or *in vitro* protein studies. We reasoned that, at least under non-reducing conditions, trypsin cleavage would result in disulfide-bound peptides between the nucleophilic YAP1C Cys598 and the sulfenylated cysteine in the target proteins. Hence, these mixed peptides entail the necessary information on the sulfenylated cysteines. Theoretically, trypsin cleavage around the disulfide bond generates a mixed peptide involving a 7-amino-acid peptide from the YAP1C probe ‘C_598_SEIWDR’ that contains the redox-active Cys598, and the sulfenylated cysteine-containing peptide from the YAP1C-bound protein (Figure 1). Typically, the strong background of the tryptic linear peptides hinders the identification of cross-linked (CL) peptides. Therefore, enrichment procedures are performed in chemical cross-link proteomics (Barysz and Malmström, 2018). Similarly, we devised an enrichment strategy for the C_598_SEIWDR CL peptides by generating polyclonal antibodies directed toward the C_598_SEIWDR peptide (see Materials and Methods). Afterward, we tested four workflows for the detection of C_598_SEIWDR CL peptides by pLink 2 (Chen et al., 2019) (Figure 1). Proteins were extracted from YAP1C cells treated with 20 mM H_2_O_2_ for 30 min. Note that for all tested workflows, equal fractions of proteins were used for LC-MS/MS (0.5 mg). Firstly, a trypsin-digested proteome was submitted for MS (shotgun proteomics), resulting in the identification of 5,917 linear (regular) peptides that provided a general proteome reference (Figure 1, sample A). However, no C_598_SEIWDR CL peptides were found, highlighting the need for YAP1C enrichment strategies. Hence, we subjected the digested proteome to an anti-C_598_SEIWDR enrichment (Figure 1, sample B). However, such direct peptide-level enrichment resulted in no C_598_SEIWDR CL peptide identifications. To reduce the peptide complexity prior to the anti-C_598_SEIWDR enrichment, the proteome was first enriched on the IgG-Sepharose beads and, second, after an on-beads trypsin digestion, the eluted sample was subjected to an anti-C_598_SEIWDR affinity purification and analyzed by MS (Figure 1, sample C). This double enrichment strategy proved highly successful, because 475 C_598_SEIWDR CL peptides were identified (≤ 1% FDR; Supplementary Dataset S1). Lastly, we tested CL peptide identifications of the eluted sample only after IgG-Sepharose enrichment (Figure 1, sample D). After protein-level enrichment, solely one single CL peptide, ‘C#SEIWDR-VIEYC#K’ (with C#, indicating a CL cysteine), was identified that matched the S PHASE KINASE-ASSOCIATED PROTEIN (SPK)-like proteins. Hence, a dedicated cross-link enrichment step is required after the IgG-Sepharose enrichment for large-scale identification of YAP1C CL peptides. Moreover, the 6,721 identified linear peptides provide complementary information on the YAP1C protein interactors. Taken together, analysis of YAP1C protein interactors, after IgG-Sepharose enrichment, followed by the identification of YAP1C CL sites, after an additional anti-C_598_SEIWDR enrichment step, enables the proteome-wide detection of sulfenylated cysteines.

**Figure 1.**
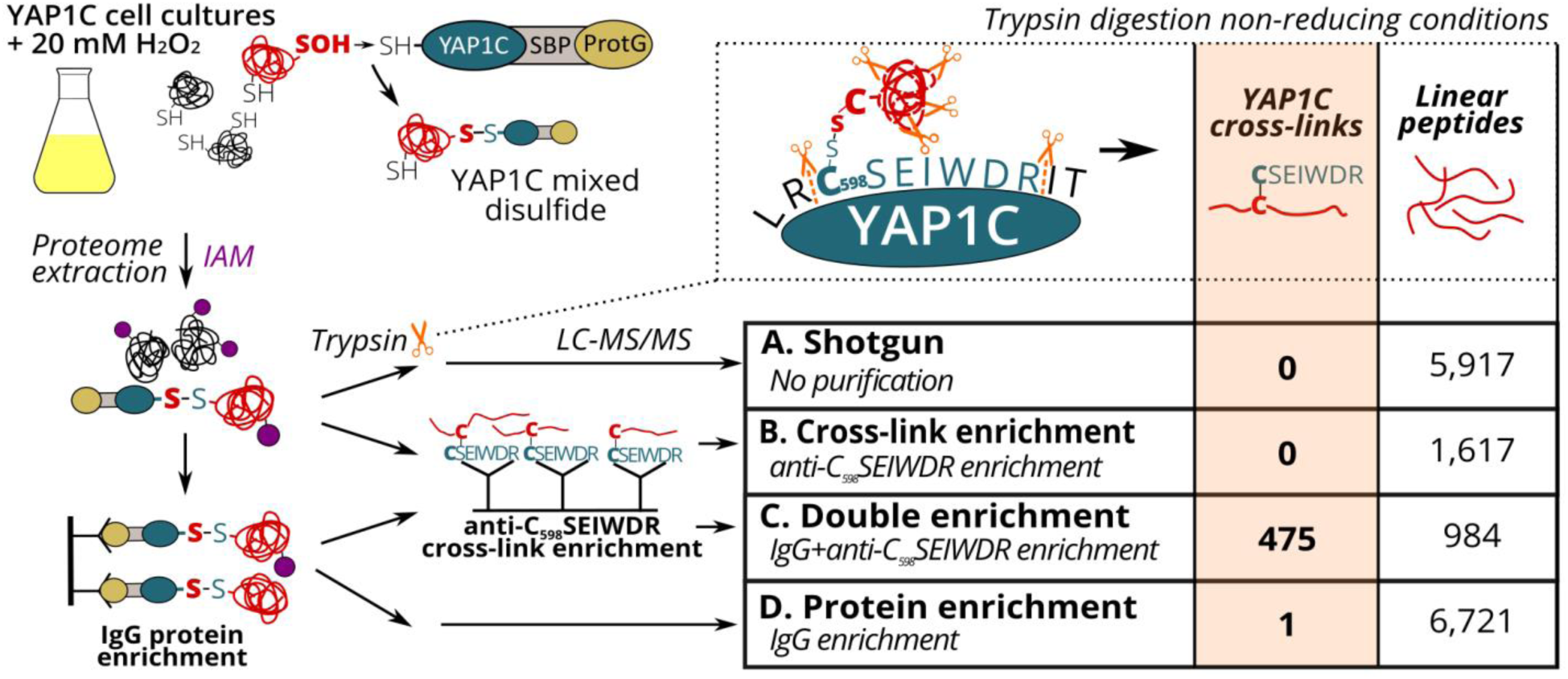
Enrichment strategies for YAP1C cross-linked (CL) peptide identification. Trypsin digestion under non-reducing conditions results in YAP1C C_598_SEIWDR CL peptides containing the redox-active Cys598. Next to a nonenriched proteomic shotgun (sample A), CL peptide enrichment was tested with anti-C_598_SEIWDR polyclonal antibodies (sample B). In addition, YAP1C-interacting protein complexes were purified on IgG-Sepharose beads, followed by an on-bead trypsin digestion (sample D). The obtained peptides were further enriched for C_598_SEIWDR CL peptides on beads coupled to the anti-C_598_SEIWDR antibody (sample C). Abbreviations: IAM, iodoacetamide; LC-MS/MS, liquid chromatography-tandem mass spectrometry; ProtG, two protein G domains; SBP, streptavidin-binding peptide.

### Identification with YAP1C of sulfenylated cysteines under H_2_O_2_ stress

To identify sulfenylated cysteines in untreated and treated (20 mM H_2_O_2_ for 30 min) YAP1C cells (three replicates per condition), we analyzed YAP1C protein complexes after IgG-Sepharose enrichment and their YAP1C CL sites after an additional anti-C_598_SEIWDR enrichment step. In total, 1,930 C_598_SEIWDR CL peptides were detected (7,040 peptide-to-spectrum matches (PSMs); Supplementary Dataset S2). Due to trypsin missed cleavages, some CL peptides indicated the same sulfenylated cysteine. For instance, ‘C#ATITPDEGR-C#SEIWDR’ and ‘C#ATITPDEGRVTEFGLK-C#SEIWDR’, both indicate *S*-sulfenylation of Cys75 in CYTOSOLIC NADP^+^-DEPENDENT ISOCITRATE DEHYDROGENASE. Removal of this redundancy (see Materials and Methods) resulted in a total of 1,747 nonredundant C_598_SEIWDR CL peptides (Supplementary Dataset S2). Of these 7,040 PSMs, 25 (0.36%) are indicative of inter-YAP1C cross-links between the redox-active Cys598, suggesting that a minor artefactual self-trapping of YAP1C is possible. In favor of high-confident identifications, we retained 1,132 out of the 1,747 C_598_SEIWDR CL peptides with at least two PSMs across the six samples (Supplementary Dataset S2). Somehow unexpectedly, more C_598_SEIWDR CL peptides were identified in untreated samples (1,082 CL peptides) than in the H_2_O_2_-treated cells (759 CL peptides). Nevertheless, the majority of the C_598_SEIWDR CL peptides (709 out of 1,132 [63%]) was identified under both conditions and 50 CL peptides were exclusively identified after H_2_O_2_ stress (Figure 2A). Possibly, the 20-mM H_2_O_2_ treatment might result in overoxidation of certain cysteines, activation of antioxidant systems, or *S*-glutathionylation. From the 1,132 C_598_SEIWDR CL peptides, 307 could not be attributed unambiguously to a unique *Arabidopsis* protein, meaning that the YAP1C CL *Arabidopsis* peptides are present in at least two different proteins. The remaining 825 CL peptides matched uniquely to 570 different proteins, implying that some *Arabidopsis* proteins contain multiple *S*-sulfenylated sites. For 94% of the 570 YAP1C CL proteins, at least one peptide was identified after IgG-Sepharose enrichment, thus complementary evidenced as a YAP1C-interacting protein. From the 307 CL peptides that match multiple *Arabidopsis* proteins, protein-level evidence was exclusively available for solely one of the possible matching proteins. For instance, the CL peptide ‘KLKEC#EK-C#SEIWDR’ represents a C_598_SEIWDR CL to either Cys124 in PROTEIN PHOSPHATASE5 (PP5, AT2G42810) or Cys487 in PROTON PUMP INTERACTOR2 (PPI2, AT3G15340) (Supplementary Dataset S2). Prior to the anti-C_598_SEIWDR purification step, 23 linear peptides (194 PSMs) were identified for PP5, whereas none for PPI2. As such, besides the complementary evidence of the YAP1C-interacting proteins, the MS analysis of the IgG-Sepharose-enriched samples are helpful for protein identification in case of ambiguous peptide-to-protein matchings.

**Figure 2.**
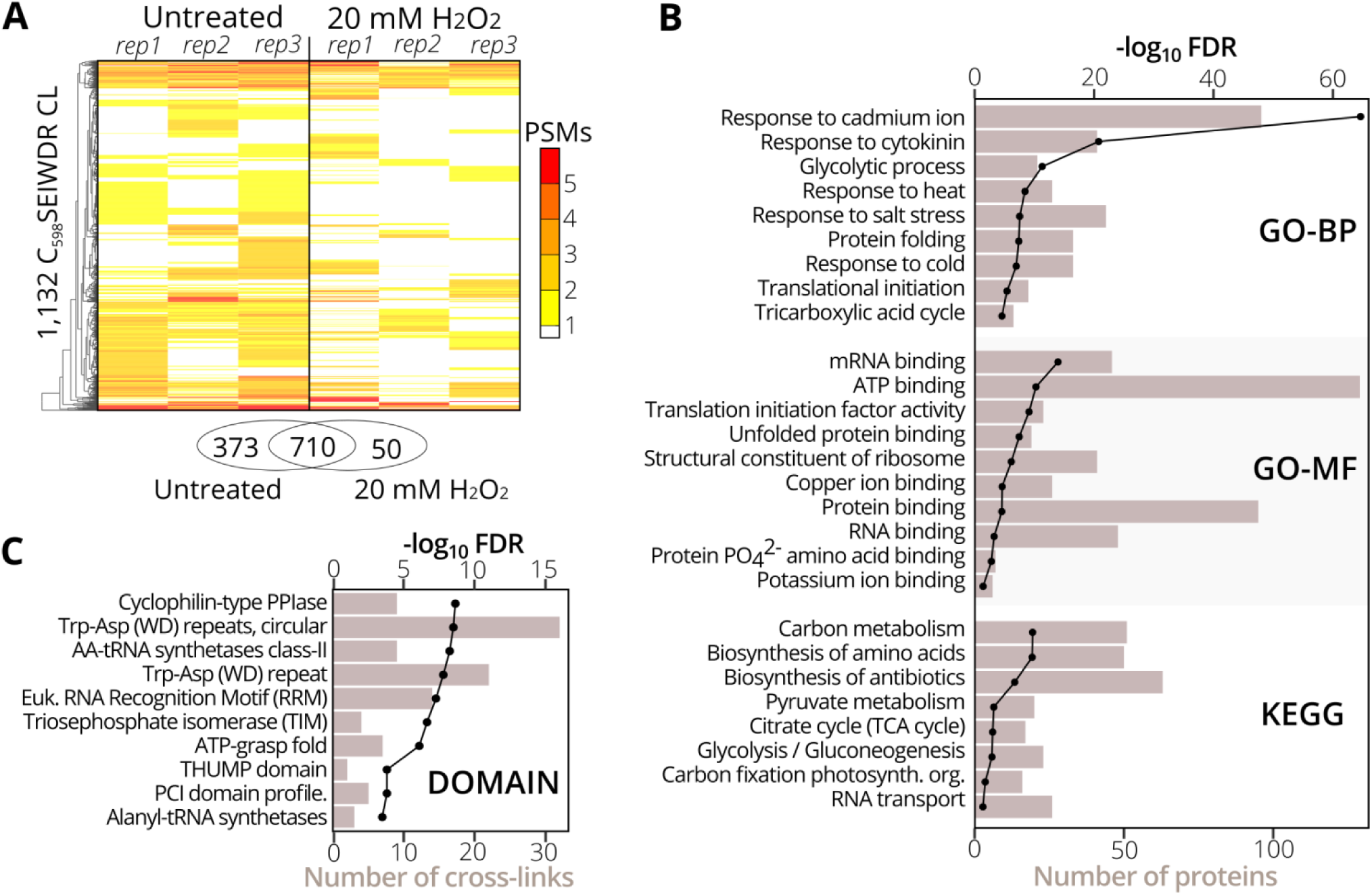
Gene set and protein domain enrichment of YAP1C C_598_SEIWDR CL proteins and sites. **(A)** Heatmap of the 1,132 C_598_SEIWDR CL peptides (FDR ≤ 1%; ≥ 2 PSMs) showing the number of PSMs per replicate in cells untreated (left) and treated with H_2_O_2_ for 30 min (right). The overlap of C_598_SEIWDR CL peptides between conditions is shown in a Venn diagram. **(B)** Gene set enrichment of the 570 proteins matched uniquely by the C_598_SEIWDR CL peptides with the DAVID tool (Huang et al., 2009). The assessed gene sets were KEGG pathways, gene ontology (GO) terms for biological process (GO-BP), molecular function (GO-MF), and cellular component (GO-CC, not plotted). All results are presented in Supplementary Dataset S3a. (**C**) Enrichment of 1,132 sulfenylated cysteines in PROSITE profiles (see Materials and Methods). Enrichment was plotted as a black line (-log_10_ FDR, top x-axis) and the number of CL sites as a bar chart (bottom x-axis). All results are reported in Supplementary Dataset S3b.

To functionally categorize the identified *S*-sulfenylated proteins, we carried out a gene set enrichment analysis on the 570 proteins for which 825 sulfenylated cysteines had unambiguously been identified by C_598_SEIWDR CL peptides. In accordance with the cytosolic localization of the YAP1C probe, the strongest overrepresented GO term was the cellular component ‘cytosol’ (FDR 2.3e^-202^; Supplementary Dataset S3a). In addition, proteins of several stress-related adaptive processes (Cd^+^, salt, heat, and cold), metabolism, and RNA processes were strongly enriched (Figure 2B), corresponding with our previous reports that demonstrated that enzymes involved in core carbon metabolic pathways, such as glycolysis, amino acid metabolism, and carbon fixation, were prone to *S*-sulfenylation (Huang et al., 2018, 2019). Also, the GO-MF term ‘mRNA-binding’ and the KEGG pathway ‘RNA transport’ were overrepresented (FDR 3.6e^-16^ and 0.04, respectively) (Figure 2B). Sulfenylated cysteines in various protein domains (Figure 2C). For instance, in the *Arabidopsis* proteome, the amino acyl-transfer RNA (AA-tRNA) synthetase domain is present in seven proteins and contains 96 cysteines in total, of which nine found within the AA-tRNA synthetase domain were detected as sulfenylated and were overrepresented (FDR 5.9e^-9^; Supplementary Dataset S3b). In addition, the RNA-binding RNA-recognition motif (RRM) was overrepresented (FDR 5.5e^-8^) that had previously been reported as a *S*-sulfenylation hotspot (Huang et al., 2019). Taken together, functional enrichment of *S*-sulfenylated proteins and cysteines identified by C_598_SEIWDR CL peptides match previous observations by other tools.

### YAP1C cross-linked peptides are associated with characteristic disulfide ions

In a next phase, we used the 7,000 PSMs of C_598_SEIWDR CL peptides to characterize in detail the general properties and MS/MS fragmentation patterns of C_598_SEIWDR CL peptides. Firstly, we compared the properties of C_598_SEIWDR CL peptides to the linear peptides identified prior to the anti-C_598_SEIWDR purification. As CL peptides are the combination of two peptides with charged N-termini and tryptic C-termini ending on Arg/Lys, CL peptides typically have both a higher mass and precursor charge than the linear peptides. In accordance, the peptide mass of C_598_SEIWDR (907 Da) approximates the median peptide mass difference of identified C_598_SEIWDR CL peptides with linear peptides (Figure 3A, 892 Da). In addition, precursors of the C_598_SEIWDR CL peptides are more positively charged than the linear peptides, with approximately 93% of the PSMs charged ≥ 3+ (Figure 3B). As such, both peptide mass and charge are in line with typical CL peptide properties. Importantly, the CL peptide identification is more challenging than that of linear peptides, because the fragmentation of CL peptides results in intermixed fragment ions derived from both peptides. Noteworthy, the CL peptide search with pLink 2 was not biased toward C_598_SEIWDR CLs and 27 intra-protein and 20 inter-protein (non-YAP1C) CL peptides were identified (102 PSMs, FDR ≤ 1%) (Supplementary Dataset S4). Hence, despite the search for CL peptides between or within 26,000 *Arabidopsis* proteins, the YAP1C C_598_SEIWDR CL peptides are by far preponderant (7,040 C_598_SEIWDR CL PSMs versus 102 non-YAP1C PSMs), and emphasize the effectiveness and necessity of the anti-C_598_SEIWDR purification. Moreover, Cys598 of YAP1C was also CL to *Arabidopsis* peptides via the trypsin-missed cleaved peptides ‘EGSLLRC#SEIWDR’ and ‘C#SEIWDRITTHPK’. For instance, the peptide ‘HMIEDDC#TDNGIPLPNVTSK’ of the E3 ubiquitin ligase SKP-like protein 1B (SKP1B; AT5G42190) was CL to both ‘C#SEIWDR’ (Figure 3C) and ‘C#SEIWDRITTHPK’ (Supplementary Dataset S2). This cytosolic protein had been recognized was identified previously to be *S*-sulfenylated by the original protein-level YAP1C-TAP strategy (Waszczak et al., 2014).

**Figure 3.**
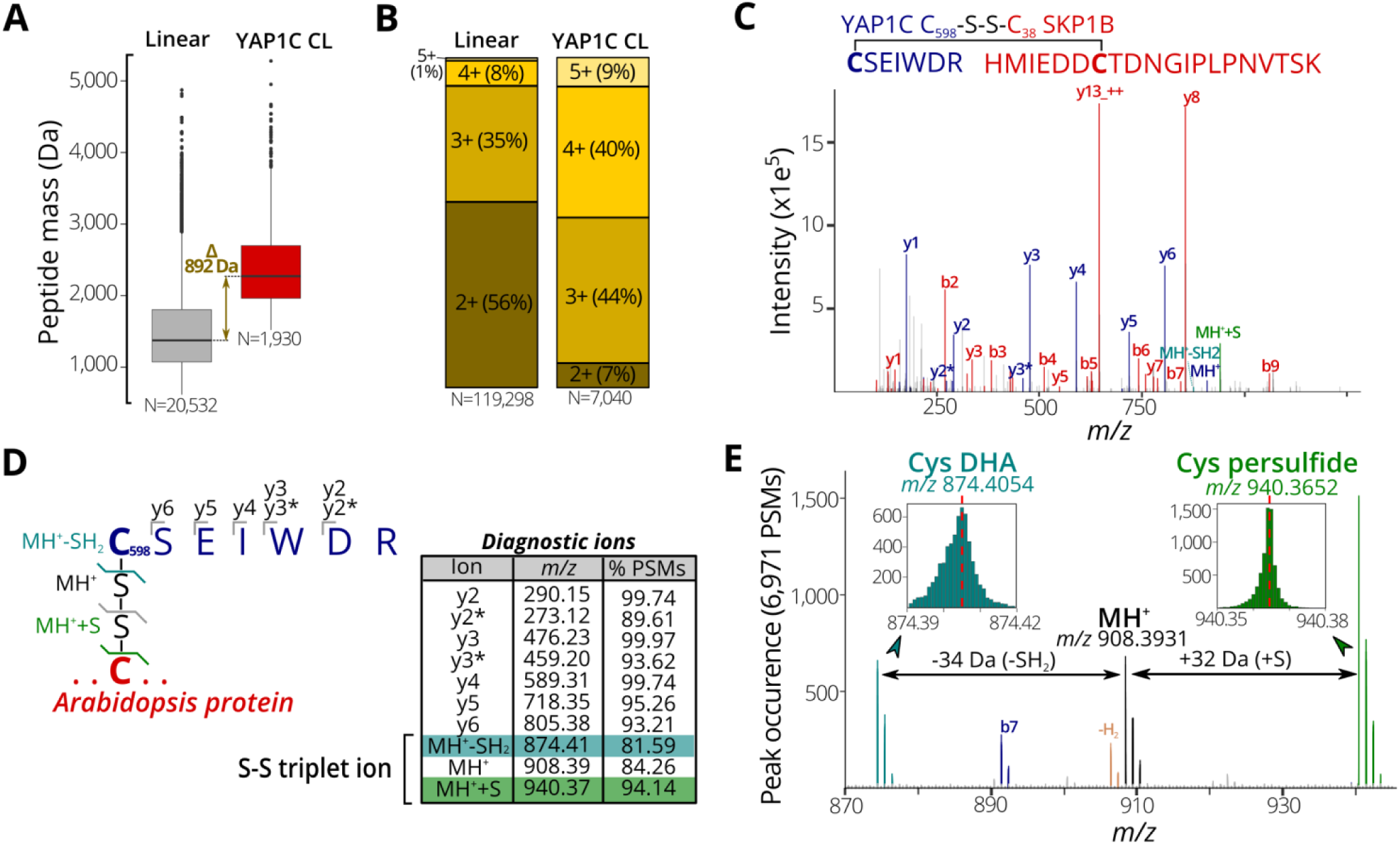
Characteristic properties and fragmentation ions of C_598_SEIWDR CL peptides. **(A)** Mass distribution for nonredundant linear and C_598_SEIWDR CL peptides (right). A 892-Da difference is indicated between the median peptide masses of both distributions. **(B)** Peptide precursor charge distribution for PSMs to linear and C_598_SEIWDR CL peptides. **(C)** Annotated spectrum of C#SEIWDR (blue fragment ions) CL to HMIEDDC#TDNGIPLPNVTSK (red fragment ions). **(D)** Parsing of 6,971 PSMs of the YAP1C cross-links (FDR ≤ 1%) containing at least five C_598_SEIWDR b-,y-, or precursor ions (within *m/z* 0.01) for diagnostic ions. To this end, the occurrence of a peak was counted at a *m/z* 0.001 interval (irrespective of its intensity) to identify consistent fragment ions. Ten characteristic C_598_SEIWDR CL peptide ions were displayed in the peptide fragmentation scheme and table, with y2* and y3* indicating an ammonia neutral loss (-NH_3_). A triplet ion resulting from disulfide fragmentation patterns is highlighted. **(E)** Occurrence of triplet ion peaks characteristic of disulfide cleavage in 6,971 PSMs of C_598_SEIWDR CL peptides (FDR ≤ 1%, ≥ 5 fragment ions). The expected C_598_SEIWDR precursor mass is indicated in black (*m/z* 908.3931) and is flanked by Cys dehydroalanine (DHA, blue; *m/z* 874.4054) and Cys persulfide (green; *m/z* 970.3652). Brown peaks correspond to the MH+ precursor minus two hydrogens corresponding to an intact disulfide bond. Abbreviations: MH^+^, C_598_SEIWDR single-charged peptide precursor; MH^+^+S, Cys persulfide precursor; MH^+^-SH_2_, Cys DHA precursor.

Next, we aimed to identify characteristic fragment ions of C_598_SEIWDR CL peptides. Such characteristic ions could help in the assessment of the PSM quality or future identification of C_598_SEIWDR CL peptides. To detect consistent fragment ions associated with C_598_SEIWDR CL peptide fragmentation, we counted the occurrence of MS/MS *m/z* peaks in 6,971 PSMs (98.8% of 7,040 total C_598_SEIWDR CL PSMs), containing at least five b-, y-, or precursor ions of C_598_SEIWDR. The C_598_SEIWDR fragment y-ions are well represented, occurring in 93% to 99% of the cases (Figure 3D), as well as y2- and y3-ions with neutral loss of NH_3_ (Figure 3D, y2* and y3*, respectively). Interestingly, besides the C_598_SEIWDR precursor ion (MH^+^, *m/z* 908.39), neighboring masses corresponding to the precursor ion with cysteine persulfide formation (blue; +S, *m/z* 940.37) and a cysteine-to-dehydroalanine conversion (green; -SH_2_, *m/z* 874.41) are consistently present in C_598_SEIWDR CL PSMs (Figure 3D and E). This distinctive pattern of triplet ions is characteristic for inter-protein disulfides (Janecki and Nemeth, 2011) and used, for instance, by dedicated disulfide CL identification algorithms, such as DBond (Na et al., 2015) and MS2DB+ (Murad et al., 2011). Together with the C_598_SEIWDR y fragment ions, these precursor triplet ions form a distinctive C_598_SEIWDR fragment ion fingerprint. For example, 4,220 PSMs (59.9%) of the identified C_598_SEIWDR CL PSMs contain all 10 of these characteristic ions, whereas 6,503 (92.4%) and 5,847 PSMs (83.1%) had eight and nine out of 10 ions, respectively. As such, these characteristic ions can help in PSM quality assessment of C_598_SEIWDR CL peptides, with, for instance, all characteristic ions present in the ‘HMIEDDC#TDNGIPLPNVTSK-C#SEIWDR’ PSM (Figure 3C). In addition, we used the characteristic fragment ions (Figure 3D) as a fingerprint to scan potential C_598_SEIWDR CL peptides in the raw proteomics data obtained after IgG-Sepharose and/or anti-C_598_SEIWDR enrichment strategies (Figure 1). After IgG-Sepharose enrichment, 33 MS/MS spectra contained the full C_598_SEIWDR CL fingerprint (Supplementary Dataset S5), indicating that more than a single C_598_SEIWDR CL might be fragmented, but not identified in the pLink 2 search (FDR ≤ 1%; Supplementary Dataset S1). Missing identifications can arise due to numerous reasons, such as the *Arabidopsis* peptide CL to C_598_SEIWDR being shorter than six amino acids (default pLink 2 search settings), noisy spectra with low-abundant fragment ions, or too stringent FDR scoring. In line with the high number of C_598_SEIWDR CL peptides identified by pLink 2 (475 peptides; Supplementary Dataset S1), 1,502 MS/MS spectra contained the full C_598_SEIWDR CL fingerprint after the additional anti-C_598_SEIWDR enrichment step (Supplementary Dataset S5). In contrast, no spectra with a full C_598_SEIWDR CL fingerprint were found in the proteome shotgun analysis or after direct anti-C_598_SEIWDR enrichment (Figure 1, samples A and B, respectively), indicating a high specificity for the fingerprint toward C_598_SEIWDR CL peptides. As such, the proposed 10 characteristic ions provide a useful and distinctive fingerprint for C_598_SEIWDR CL peptides for quality assessment of individual spectral matches or raw proteomics data.

### YAP1C cross-linked cysteines report protein redox-sensitive cysteine sites

We assessed whether the identified C_598_SEIWDR CL peptides (Supplementary Dataset S2) are in agreement with related redox studies. First, in 67 out of the 97 *S*-sulfenylated proteins (69%) identified previously as YAP1C interactors (Waszczak et al., 2014), 102 *S*-sulfenylated sites were found, including, for instance, Cys20 of DEHYDROASCORBATE REDUCTASE2 (DHAR2) (Figure 4A) that had been shown to be *S*-glutathionylated via an *S*-sulfenylation intermediary by 5,5-dimethyl-1,3-cyclohexadione (dimedone) labeling and MS identification of recombinantly produced DHAR2 (Waszczak et al., 2014; Bodra et al., 2017). This laborious approach to identify the trapped *S*-sulfenylation sites can be avoided thanks to the identification of C_598_SEIWDR CL peptides, as shown here for DHAR2. Taken together, this procedure will help hypothetically formulate the mode-of-action of potential redox switches and fast-forward downstream experiments.

**Figure 4.**
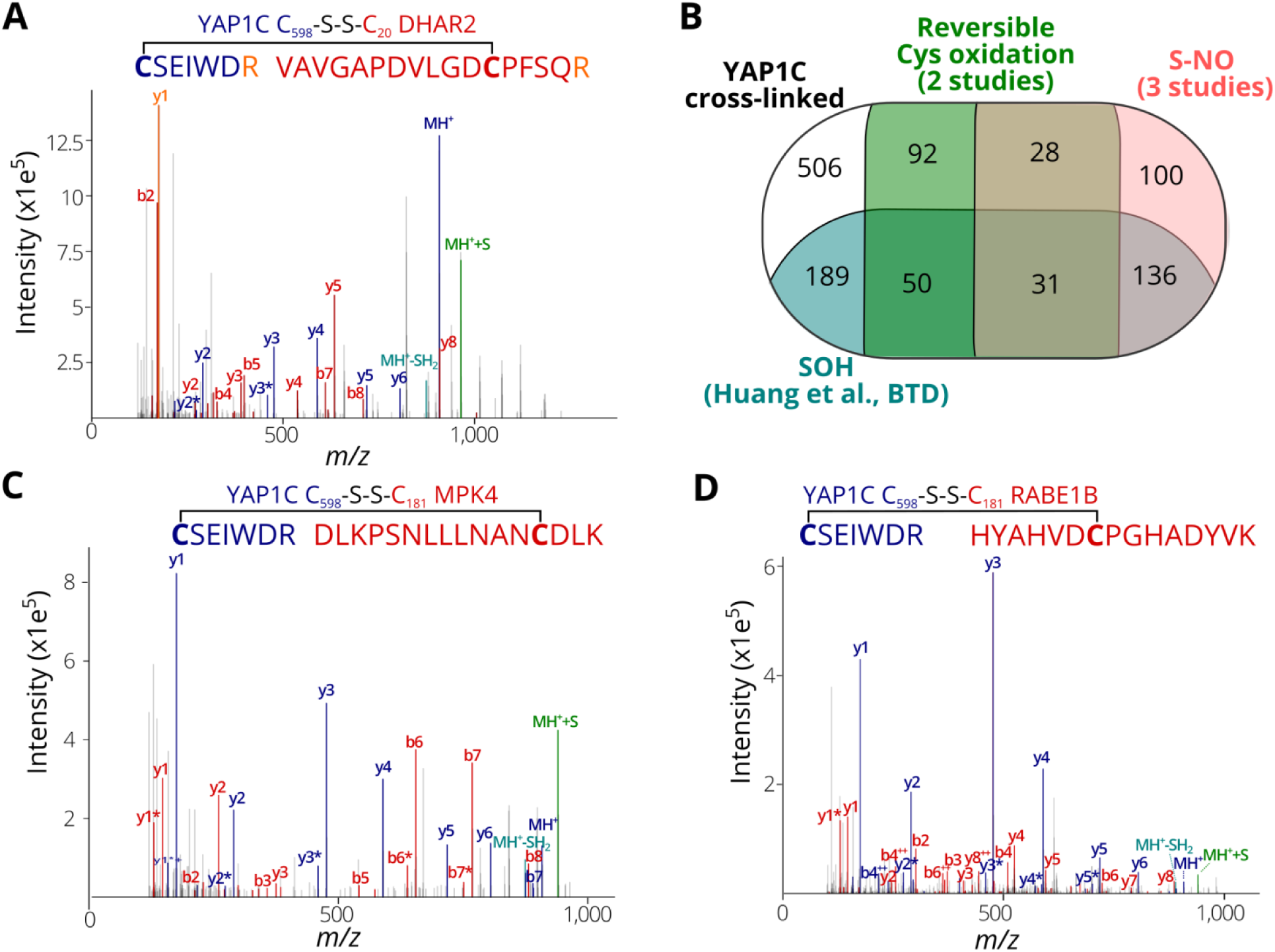
*S*-sulfenylated sites identified by C_598_SEIWDR CL peptides corresponding to previously identified redox-sensitive cysteines. **(A)** Annotated MS/MS spectrum of C#SEIWDR (blue) linked to VAVGAPDVLGDC#PFSQR (red, Cys20 of DHAR2). The y1 fragment ion (corresponding to the C-terminal Arg) (orange) is shared by both peptides. **(B)** Venn diagram displaying the overlap of 1,132 *S*-sulfenylated sites (≥ 2PSMs) identified in this study (black) with previously reported *Arabidopsis S*-sulfenylated sites (blue) (Huang et al., 2019), *S*-nitrosylated sites (pink) (Fares et al., 2011; Puyaubert et al., 2014; Hu et al., 2015) and reversibly oxidized cysteine sites (green) (Liu et al., 2014, 2015). **(C)** Annotated MS/MS spectrum of C#SEIWDR (blue) linked to DLKPSNLLLNANC#DLK (red; Cys181 of MPK 4). **(D)** Annotated MS/MS spectrum of C#SEIWDR (blue) linked to VAVGAPDVLGDCPFSQR (red). For all annotated MS/MS spectra, the C_598_SEIWDR fragment ions are colored in dark blue. Persulfide and cysteine-to-dehydroalanine fragment ions are in green and light blue, respectively. Fragment ions of *Arabidopsis* peptides linked with sulfenylated cysteines are in red.

Then, the identified *S-*sulfenylated cysteines by C_598_SEIWDR CL peptides were compared to redox-sensitive *Arabidopsis* cysteines reported previously. Recently, we have identified more than 1,500 sulfenylated Cys residues in *Arabidopsis* cells by means of a chemoselective benzothiazindioxide (BTD)-based carbon nucleophile (Huang et al., 2019). Comparison of the 1,132 YAP1C CL peptides (Supplementary Dataset S2; ≥ 2 PSMs) revealed 406 sites (36%) that overlapped with BTD-targeted Cys (Figure 4B). For instance, the C_598_SEIWDR CL peptides ‘C#SEIWDR-DLKPSNLLLNANC#DLK’ (Figure 4C) matched the Cys181 of MITOGEN-ACTIVATED PROTEIN KINASE4 (MPK4) that had been experimentally verified (Huang et al., 2019). Also, other site-specific reversibly oxidized cysteine studies (Liu et al., 2014, 2015) and *S*-nitrosylation studies (Fares et al., 2011; Puyaubert et al., 2014; Hu et al., 2015) were compared (Willems et al., 2019). In total, 295 *S*-nitrosylation and 201 reversible previously reported cysteine oxidation sites overlapped with the YAP1C-CL sites (Figure 4C). Taken together, 626 out of the 1,132 sulfenylated cysteines identified here by C_598_SEIWDR CL peptides (55.3%) had already been reported as redox sensitive in independent studies, of which some had been confirmed biochemically. For instance, ‘C#SEIWDR-HYAHVDC#PGHADYVK’ matched Cys149 of the chloroplastic elongation factor Tu (EF-Tu) RAB GTPASE HOMOLOG E1B (RABE1B), a site identified previously as *S*-sulfenylated (Huang et al., 2019) and *S*-nitrosylated (Hu et al., 2015). Interestingly, sulfenylation of the corresponding cysteine site (Cys82; Supplemental Figure S1) in the EF-Tu ortholog of *Cyanobacterium synechocystis*, the chloroplast cyanobacterial ancestor, inactivates EF-Tu in a reversible manner (Puyaubert et al., 2014; Yutthanasirikul et al., 2016). Taken together with the approximately 3.5-fold increased *S*-sulfenylation under H_2_O_2_ treatment (Huang et al., 2019), the plastidial EF-Tu protein probably exhibits a conserved redox sensitivity in *Arabidopsis*. All in all, the high agreement of the *S*-sulfenylated sites identified by C_598_SEIWDR CL peptides with other studies of redox-sensitive cysteines demonstrates that the YAP1C-resolved cysteines are in general highly susceptible to oxidative redox modifications. Identification of these redox-sensitive sites will greatly facilitate the possible formulation and our understanding of the redox signaling processes.

## CONCLUSION

Here, we report an innovative approach for *in situ* identification of *S*-sulfenylation sites by means of a transgenic probe YAP1C. By IgG purification at the protein level, followed by anti-C_598_SEIWDR purification at the peptide level, large-scale capture and identification of YAP1C CL peptides are possible, thereby uncovering *in vivo S*-sulfenylated protein sites. Importantly, this method can detect sulfenylated cysteines in a noninvasive manner and might easily be adapted to detect sulfenylated cysteines in specific organelles (De Smet et al., 2019). The proposed genetically based methodology holds great promise for *in planta* mining of *S*-sulfenylated sites, in which rigid plant tissues limit the penetration and use of chemoselective probes. All in all, thanks to this noninvasive approach based on the YAP1C probe, the site-specific identification of protein *S*-sulfenylation was successfully shown for *Arabidopsis* cells under H_2_O_2_ stress. Importantly, this method can be implemented for any species that can be genetically modified.

## AUTHOR CONTRIBUTIONS

B.W., P.W., J.H., J.M. and F.V.B. conceived the research; B.W., C.T.; and J.Y. conducted the experiments; B.W. and P.W. analyzed the data; and B.W., P.W., J.H., J.M., and F.V.B. wrote the paper.

## Supporting information

Cross-linked peptide report by pLink 2 of purification test samples

Integrative overview of 1,747 YAP1C-cross-linked peptides

gene set enrichment analysis

Integrative overview of 47 non-YAP1C cross-linked peptides

MS/MS spectra containing the ten characteristic CSEIWDR CL peptide fragment ions

## ACKNOWLEDGMENTS

The authors thank Dr. Didier Vertommen and Dr. Barbara De Smet for helpful discussions, Dr. Lieven Sterck for submitting data to the Plant PTM Viewer, and Dr. Martine De Cock for help in preparing the manuscript.

## SUPPLEMENTARY MATERIAL

**Supplementary Figure S1.**
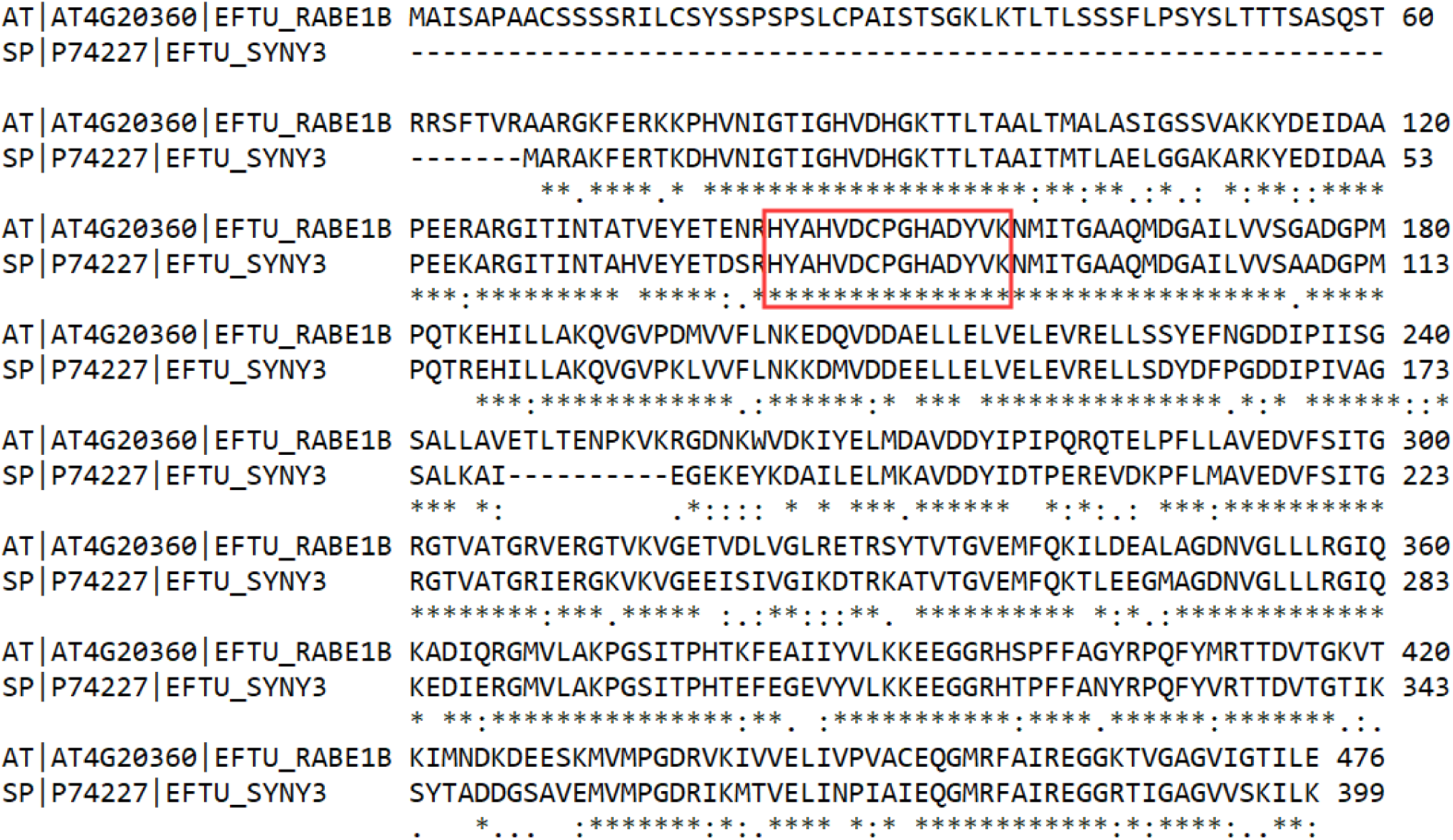
Protein alignment between EF-Tu proteins of *Cyanobacterium Synechocystis* (SP | P74227 | EFTU_SYNY3) and Arabidopsis (AT | AT4G20360 | EFTU_RABE1B). The peptide in red box was identified as YAP1C cross-linked in this study (Supplementary Dataset S2).

